# Spatial capture-recapture reveals resource use and declining skipper numbers in Baltic Sea salmon trolling fisheries

**DOI:** 10.1101/2025.01.13.631546

**Authors:** Konrad Karlsson

**Affiliations:** Department of Aquatic Resources, Institute of Freshwater Research, Swedish University of Agricultural Sciences, Stångholmsvägen 2, 178 93 Drottningholm, Sweden

**Keywords:** Baltic Sea, fish conservation, fisheries management, recreational fisheries, Atlantic salmon, spatial ecology

## Abstract

Many populations of Atlantic salmon (*Salmo salar*) native to the Baltic Sea are endangered. A significant group targeting salmon in this region is the recreational trolling fisheries. Due to frequent changes in fisheries regulations and conditions, both the conservation of the species and the fishers are impacted. This study aims to enhance our understanding of the spatial extent and population dynamics of salmon trolling boat skippers in the Baltic Sea by estimating their population size and resource utilization. The study utilizes participation lists of skippers from recreational fishing tournaments and Automatic Identification System (AIS) data to track their movements. These lists are formatted as encounter histories for spatial capture-recapture models, while AIS data are integrated as telemetry data to estimate resource selection. The results reveal a 51% decrease in the number of skippers from two time periods: years 2014-2020 and 2021-2023, with the count dropping from 5,343 individuals (95% CI: 4,622-6,178) to 2,604 individuals (95% CI: 2,273-2,983). The decline may be attributed in part to heavy regulations imposed on recreational salmon fisheries in 2022. Furthermore, the resource selection analysis indicates that these skippers target various species also outside of the Baltic Sea, such as Bluefin tuna (*Thunnus thynnus*) in Skagerrak, and endangered salmon stocks elsewhere, such as in Lake Vänern. The results of this study suggests that regulations and changes in the Baltic Sea salmon trolling fishery may have broader impacts on seemingly unrelated species and ecosystems.

## Introduction

Effective management of fisheries requires an understanding of resource selection and population dynamics. The Atlantic salmon (*Salmo salar*) is a keystone species in both recreational and commercial fisheries, as well as in aquaculture. However, many salmon populations are in decline, endangered, or extinct due to habitat loss and other anthropogenic pressures (Klemetsen et al., 2003; Kesler et al., 2011). In the Baltic Sea, the primary driver of declining salmon numbers is the loss of riverine habitats, with hydropower development contributing significantly to population extinctions (HELCOM, 2011).

Salmon in the Baltic Sea are confined to its basins and do not extend through the Danish straits (Kallio-Nyberg and Ikonen, 1992). The region hosts at least 43 self-reproducing populations native to distinct river systems, so called wild salmon. Although many populations are maintained through brood stocking programs due to habitat loss (HELCOM, 2011). These stocked salmon, often marked by adipose fin removal, enable selective fishing pressure between farmed and wild fish (HELCOM, 2011). Adult salmon mix in southern and southwestern Baltic feeding grounds before returning to their natal rivers to spawn, while younger fish remain closer to home (Palm et al., 2008; Whitlock et al., 2018; Jacobson et al., 2020). Because small and large populations mix this complicates efforts to protect vulnerable populations, such as the small population in Estonia’s Purtse River, alongside larger populations in Kalix and Torne/Tornio rivers in Sweden and Finland (Kesler et al., 2011; Kärgenberg et al., 2019; Miettinen et al., 2021).

To safeguard small and vulnerable populations, a series of restrictions has been implemented in the open sea. Within recreational sea fisheries, salmon landings are predominantly made by trolling fisheries (ICES, 2021; Hartill et al., 2020). Therefore, when regulating the sea-based salmon fishery of the Baltic Sea, trolling fisheries has been a key focus.

Baltic Sea salmon stocks native to specific rivers may traverse several nations’ borders while at sea. Just a few years ago the Baltic Sea salmon trolling fishery was regulated by a complex mix of international, national, and regional harvest policies. Recent restrictions aim to streamline policies and protect vulnerable populations, notably Council Regulation (EU) 2023/2638, which prohibits wild salmon landings in Baltic Sea areas 22–31 while allowing limited recreational fishing of marked (farmed) salmon. Thus, prior to 2022, there were significant regional disparities and more generous harvest regulations, which have now been streamlined due to EU legislation (ICES, 2021; ICES, 2023). However, some exemptions and regional disparities remain, such as national regulations in northern areas, where fishing restrictions differ by latitude.

For example, in Sweden (FIFS 2004:36), from May 1st to August 31st, within 4 nautical miles from the baseline north of latitude 63°30′N harvest of wild salmon is prohibited, meanwhile the number of harvestable farmed salmon during recreational trolling is unlimited. During the years 2022-2023 this legislation applied up to 59°30′N (EU 2022/2090). It was later moved northwards to protect further populations in the south, as the northernmost Baltic salmon populations are considered the most viable (ICES, 2023). These new and stringent regulations may influence the trolling fleet, a key question this study aims to address.

Understanding the dynamics of the trolling fleet and the impact of regulatory changes is essential to estimate the number of skippers operating recreational salmon trolling boats in the Baltic Sea. Two major salmon fishing tournaments, the Trolling Master Bornholm and the Trollingträff Åland, provide annual participation lists of skippers. These lists, combined with capture-recapture modeling, enable population size estimation.

Capture-recapture models were initially developed to monitor animal populations through marking and recapture (Otis, 1978) and have since been extended to human populations using lists of individuals instead of physical captures (Chao et al., 2001). Capture-recapture models estimate population size by predicting the number unobserved individuals based on the number of observed individuals and their detection probability. While the first developed primitive detection models assumed homogeneous detection probabilities across all dimensions (Otis, 1978), current models are developed to handle the real-world scenarios which often involve heterogeneity due to individual behavior, temporal variation, and spatial factors (Royle et al., 2013a). Modern capture-recapture models have evolved to account for these complexities, incorporating covariates and advanced frameworks to address such heterogeneity (Royle et al., 2018). For example, behavioral covariates account for the positive or negative effects of previous detection on future detection probabilities.

This has particular relevance to the trolling fleet, where skippers who enjoy participating in tournaments are more likely to be detected, while non-participating skippers have lower detection probabilities. Ignoring a positive behavioral response can bias estimates by overestimating detection probability and underestimating population size, while the opposite is true for a negative response (Borchers and Efford, 2008; Chapter 7 in Royle et al., 2013a). Incorporating behavioral covariates corrects for this detection heterogeneity and improves the population size estimate.

Spatial capture-recapture (SCR) models build on traditional methods by incorporating spatial information to model movements, habitat use, and resource selection (Royle et al., 2013b; Fuller et al., 2016). A key advantage of SCR is its ability to explicitly link population size to geographic area, enabling density estimation while accounting for spatial biases in detection (Efford, 2004). Detection probabilities in SCR are modeled as a function of the distance between an individual’s home range center and a detection device, improving accuracy compared to non-spatial models. SCR also uses spatial recaptures to estimate individual home range centers and movement parameters, which are formally linked to the model’s detection probability function and to the population’s spatial extent (Efford, 2004; Sutherland et al., 2019).

Telemetry data, such as automatic identification system (AIS) data from trolling boats, can further enhance SCR models by providing high-resolution information on individual movements. This assumes that telemetered individuals behave similarly to the rest of the population (Linden et al., 2018). By integrating telemetry data, SCR models reduce the need for extensive spatial sampling and improve estimates of spatial use and resource selection (Linden et al., 2018). Telemetry data can also be used to develop resource selection functions (RSFs), which describe the relative importance of different habitats or fishing grounds.

This study combines participation lists (SCR data) with AIS telemetry data to estimate the number of trolling boat skippers from 2014 to 2023 and assess changes in response to recent regulations. By modeling skipper movements and resource selection, the study identifies fishing grounds’ relative importance and explores how restrictions might shift skipper activity. Such shifts could impact populations beyond those targeted by the regulations, underscoring the need for spatially informed management strategies.

## Methods

### Adherence to the general data protection regulation (GDPR)

Personal data were collected with as little information about the participants as possible. The tournament participation lists used in this study are public and were available from the organizers’ webpages. The Trolling Master Bornholm (TMB) is broadcast on the regional Danish television channel TV2/Bornholm, and the Trollingträff Åland (TTÅ) takes place at Käringsund Resort & Conference on the island of Eckerö in the Åland archipelago. Although participants are aware that the personal information they provide to the organizers is public, they do not consent to the data being part of the type of study presented here. Therefore, the data presented within and attached to this study has been anonymized. Furthermore, consent has not been requested since the number of people involved is large (> 1,000), their contact information is incomplete, and this is a scientific study, which reduces the need for informed consent according to Chapters 11 and 14 of the GDPR. This issue was discussed with the university’s data protection support.

### Data collection: Creating a data frame of individual encounter histories based on tournament data

The southwestern and western parts of the Baltic Sea are where the majority of salmon feed (Kallio-Nyberg and Ikonen, 1992; Jacobson et al., 2020). When the salmon return from their feeding grounds to their great Northern Rivers, they mainly use the western and central parts of the Baltic Sea as their main migration route. As they progress north, they pass through the Åland Archipelago and along the eastern shores of the Bothnian Sea (Siira et al., 2009; Jones et al., 2022). Consequently, popular trolling areas include off the Danish island of Bornholm, which is situated in the center of the salmon’s southern feeding ground. Another important area is between the Swedish mainland and the Finnish island of Åland which the salmon pass when migrating north. At these locations two major salmon trolling tournaments, The Trolling Master Bornholm (TMB) and The Trollingträff Åland (TTÅ), are held annually.

Encounter histories, i.e., lists of participants from the trolling tournament TMB, were sourced from the web, and lists from TTÅ were obtained upon request after contacting the tournament organizers. The trolling tournaments are held annually, and data were collected for the years 2014-2023. During the COVID-19 pandemic, the TMB tournament was canceled in 2020 and 2021, while TTÅ still had participant lists for those years. For the years 2014-2017 no data were included from TTÅ. Occasions that lacks data are noted within the models trap operation file to avoid confounding no data as no detections (Sutherland et al., 2019). The TMB tournament takes place at the end of April each year, and the TTÅ begins in early June, making it possible to participate in both tournaments in a single year. The encounter data was stratified into two sessions to account for changes in the population size of trolling boats between strata, with the years 2014-2020 comprising one stratum and 2021-2023 the other. Initially the data was intended to be symmetrical and include two strata of three years each (2018-2020 and 2021-2023), but as more data allows fitting more complex models allowing for estimation of heterogeneous detection probability (see two sections below), additional years were included. Because the automatic identification system (AIS) data were not available before 2019 (see next section below) and the importance of estimating a recent population size, data were divided in stratum of seven and three years. By including strata-specific covariates for density and detection probability, these estimates can be calculated independently for each stratum, i.e. 2014-2020 and 2021-2023. Within each stratum the population is assumed to be closed, i.e. no birth, immigration, death, and emigration.

For the participation lists from TMB, information included the team name, the skipper’s first and last name, the skipper’s residence (hometown and country), and the tournament year. Similarly, for the TTÅ, the lists comprised the team name, the skipper’s first and last name, the skipper’s residence (not hometown but country), and the tournament year. None of this personal data is displayed in the following results or in the attached data of the study.

To create unique identifiers for encounter histories, the skipper’s names were utilized. Thanks to the combination of the aforementioned information, it was possible to create highly credible unique identifiers for the skippers. However, because there is no conclusive way to identify individuals without a personal identification number, this data contains some degree of measurement error.

Although this is the case in nearly all real-world field data (Kéry and Royle, 2021), it is to some extent more apparent in this dataset. These data are mostly prone to false positive errors arising from participants writing their names differently from tournament to tournament. In contrast, false negatives are less likely but may occur if two different skippers share the same name and cannot be distinguished using auxiliary information.

In order to create the: participation, by tournament location, by year data frame needed to fit the spatial capture-recapture models (SCR), all yearly tournament data were stacked by row. In other words, the participation correspond to the individual detections (*i*), the tournament to the geographic location of the trap (*j*), and the year to the occasion (*k*) the individuals were detected. This three-dimensional data (*ijk*) is the core of the SCR encounter probability model (Royle et al., 2018). In total, there were 2753 entries with skipper names from TMB and 394 from TTÅ.

Names were occasionally inconsistently formatted across tournaments, with variations including the inclusion or omission of middle names, different orders of first and last names, misspellings, or differing abbreviations. In these instances, the skipper’s hometown, team name, tournament year, and country were used to identify discrepancies. If these combinations suggested that it was the same or a different skipper, the skipper’s name was edited to ensure consistency or discrepancy across years and tournaments.

To detect these formatting issues, several steps were taken. The first was to transform all names to lowercase letters. Then, a character distance matrix was created using the stringdistmatrix function in the stringdist package to calculate the Levenshtein distance matrix. A distance of 0 indicates that names are identical, and a distance of 1 indicates that one character is different, and so on. Names with one-letter differences were corrected. Second, all special characters and non-English letters (e.g., ü, å, ä, ö, ø) were replaced with English letters (u, a, a, o, o), and all double and triple spaces were replaced with a single space.

This was followed by splitting the name strings at every single space (“”), and then concatenating the first and last names, creating a new variable omitting middle names. After that, the Levenshtein distance matrix was repeated, and names with differences of up to three letters were scrutinized and corrected. The next step was to check for and correct identical matches between first names and last names, which indicates if participants reversed their order when signing up for the tournaments. This was followed by using the ‘nchar’ function to count the number of characters in the names to detect the use of initials.

Skippers can only participate once per tournament and year. Hence, by looking for duplicated entries of the same name, distinct individuals with the same name could be detected and corrected. A further check was to split the data frame by team name and browse through the skipper names of every team. This detected a few skippers that had used last and middle names interchangeably and used nicknames. As a final check that the encounter data were correct, the number of occurrences of a unique name per year was checked to determine whether it was 1 or 2, depending on the availability of data from one or two tournaments. The maximum possible occurrences were 1 in 2014, 2015, 2016, and 2017, as only TMB data were sourced, and 2 in 2018, 2019, 2022, and 2023, when both TMB and TTÅ data were available. In 2020 and 2021, the COVID-19 pandemic led to the cancellation of TMB, leaving only TTÅ, which meant a maximum of 1 occurrence was possible in those years. Consequently, there could be a total of 8 occurrences in TMB and 6 in TTÅ across all years, and these totals were confirmed.

### Automatic identification system data – used as telemetry data in the spatial capture-recapture model

Automatic Identification System (AIS) data were obtained from Sjöfartsverket (the Swedish Maritime Administration). The data included the position of maritime traffic for every day in 2019 and 2022 for the whole of Sweden and adjacent nations. AIS transponders are not mandatory for ships with a gross tonnage of less than 300, which excludes essentially all trolling boats. When installed on non-mandatory vessels, these AIS transponders are typically class-B transponders, which are designed for smaller vessels and have lower transmission power and functionality compared to the class-A transponders used on larger, mandatory vessels. This means only vessels of skippers that chose to install a transponder can be found in this dataset. Additionally, data are only transmitted if the skipper chose to use the transponder, as the skipper can choose when to switch the system on or off. (AIS transponders are governed by international regulations set by the International Maritime Organization (IMO), and more information can be found on the organization’s webpage.)

Data were first filtered to select class-B transponders and then for pleasure crafts and fishing ships. The AIS data included information about the ship name, country, time and date, and a maritime mobile service identity (MMSI) that is unique for the vessel. In the AIS data, the ship names were concatenated with the respective country and run for matches in the tournament data where the team names were concatenated with the respective country. Hence, ship names in the AIS data often matched with team names in tournaments. This resulted in 116 and 187 exact matches, respectively, for the years 2019 and 2022, where there was only one unique vessel with this name and country according to the MMSI. These ships were retained from the data. Respectively, seven and four of these ships were, according to the tournament data, operated by more than one skipper and were therefore removed. Additionally, there were two and six skippers, respectively per year, who had operated two ships each, and since the capture-recapture data is linked to skipper and not ship per se, the AIS data from these ships were merged by skipper name.

The AIS telemetry fixes were incorporated in the spatial capture recapture models to estimate resource selection and the movement parameter σ (see section below). The AIS contains somewhat irregular registrations but commonly in a few seconds’ intervals. Telemetry fixes in spatial capture-recapture are assumed to be independent observations of the position of the individual. Although complete independence is unlikely (Royle et al., 2013a), a way to mitigate this independence is to thin the observations. As a first step, this was achieved by creating hourly fixes by calculating the mean position within every hour per skipper. As a next step, skippers with <50 observations were removed, resulting in 24 and 139 remaining skippers. Fifty fixes were sampled at random to be used as telemetry fixes in the resource selection function implemented by the oSCR package (Sutherland et al., 2019). Sampling was done using a fixed random number generator so that results would be identical each time the code was executed. The 2019 and 2022 telemetry data were matched with the tournament detections of 2014-2020 and 2021-2023, respectively; this resulted in 23 and 111 individuals in each telemetry data frame.

### Density estimation and creating the state space and resource selection data frame

Spatial capture-recapture (SCR) models have a hierarchical structure that combines two key components: an observation model that describes the detection process of individuals at specific spatial locations, and a latent ecological process model that estimates the spatial distribution, movement, and abundance of the population (Sutherland et al., 2019). The standard SCR model was used to estimate detection probability. This model assumes that the detections of individuals are random Bernoulli events, *y*_*i*_ ∼ Bernoulli(*p*) for *i* = 1,…, *N*. Further, the model assumes that detection decreases with Euclidean distance from the individual’s home range center *s*_*i*_ to the trap location *x*_j_ (Efford, 2004). In this study, the effective trap locations are the locations of the trolling tournaments, and the home range centers are latent variables to be estimated. The decreasing detection probability is described by the ‘half normal’ model as Pr(*y_ijk_* = 1)= p_0_ exp[−α_1_*distance*(*x*_j_, *s*_*i*_)^2]^, where *i* is the individual, j is the trap location, and *k* is the sampling occasion. p_0_ is the intercept, which corresponds to the capture probability of an individual at the center of its home range, that is, where the distance between the trap and the home range center is zero, *distance* (*x*_j_, *s*_*i*_)^2,^ = 0, (Efford, 2004). The covariate α_1_= 1⁄2σ^2^_*i*_, where σ is the movement parameter of the half normal detection function and describes the decay rate of the detection probability with increasing distance.

Because different subpopulations may have different detection probabilities (Borchers and Efford, 2008), the detection model included four covariates to accommodate this. A behavioral response estimates whether individuals previously detected have a different detection probability. This is important to include if there is a group of highly devoted tournament skippers that have a greater detection probability than the baseline consisting of skippers that have not yet participated in a tournament. A session covariate that stratifies the population and allows for different detection probabilities per time period, 2014-2020 and 2021-2023. A binary covariate, allowing skippers residing in Denmark to have a different detection probability than skippers residing in other nations. The rationale for this was the large number of skippers with Danish residence in the data, and the short distance from Denmark to the most popular fishing grounds. Finally an area covariate was included to estimate differences in detection probability depending on a set of predetermined areas that are believed to be important for trolling anglers, described further below. Using a complementary log-log link, the detection probability (p_0*i*j*i*_) is modeled as:

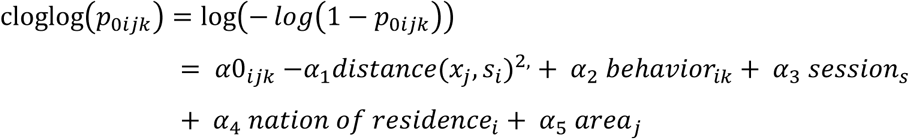

where α_0_,…, α_5_ are parameters to be estimated.

The same binary national residency covariate was included for the σ model allowing for different movement parameters for Danish and non-Danish residents. The sigma model can be expressed as:

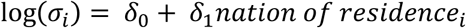

The state space ***S***, which is the spatial scale where density is estimated, had a resolution of 50 x 50 km. Based on telemetry fixes taken from the automatic identification system (AIS) data, the state space resolution corresponds to approximately half of the mean maximum distance moved (MMDM) * 0.5. This value typically lies close to the modeled σ estimate. MMDM was exp(10.7) for 2019, and exp(11.6) for 2022. A state space resolution of 0.5σ is a rule of thumb of the half-normal encounter model (Sutherland et al., 2019). The state space should include the activity centers *s*_*i*_ of the skippers that were detected in the tournaments, and its area is therefore based on the distance moved by the individuals of the population (Sutherland et al., 2019). The state space buffer was defined as the area extending 400 km from the two tournament locations which corresponds to about 3 to 4σ. At this distance detection probability is essentially zero and is therefore used as the state-space border.

Home ranges were assumed to be uniformly distributed across the state space pixels *g*, but their expected density was allowed to differ between sessions accommodated by stratifying the population per time period 2014-2020 and 2021-2023:

*log(E(D_g_)) = β_0_ + βsession_s_*

Pixel-specific frequencies from the telemetry data were used to model multinomial probabilities of space use *m*_*ig*_ ∼ Multinomial(*R*_*ig*_, *π*_*ig*_), where *R*_*ig*_ is the number of telemetry fixes for individual *i* in pixel *g*, and *π*_*ig*_ is the relative probability of use for individual *ii* in pixel *g* (Royle et al., 2013b; Linden et al., 2018), such that:

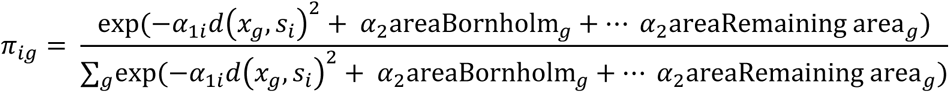

here only two of the total six areas are shown to simplify the equation above. The resource selection covariate was created by calculating the distance from the state space pixels to five distinct points representing five areas that are popular among troll anglers: Bornholm, Danish straits, Vänern and Vättern, Skagerrak, and Åland. If the distance was ≤100 km, the state space pixel was assigned the area factor. The remaining pixels that are >100 km from these five points were assigned simply as the factor level ‘remaining area’.

By integrating the joint likelihood function of telemetry data and spatial trap captures in the SCR model, the spatial resolution of the movements of individuals can be estimated both by trap detections and telemetry fixes (Royle et al., 2013a; 2013b; Linden et al., 2018). Telemetry data also allows for estimating resource selection, by modelling pixel specific relative probability of use, which reveals if certain pixels of the state space are more important than other for the population (Linden et al., 2018). Pixels with a greater capture probability have a greater importance for resource use by the individuals of the population. Linden et al. (2018) modified the integrated RSF-SCR likelihood (Royle et al., 2013b) to accommodate the lack of independence between data sources. This is useful when individuals in the SCR data are available in the telemetry data. This dependence can be modeled in the oSCR package (Sutherland et al., 2019).

Together with null models including all covariates resulted in 64 unique model combinations, models were ranked after AIC to select the best one.

## Results

Fifty hourly fixes per skipper from the Automatic Identification System (AIS) positions were randomly sampled from 24 skippers in 2019, and 139 in 2022 (Fig. 1ab). Of these, 23 and 111 skippers were detected in the tournament lists during the corresponding sessions (2014-2020 or 2021-2023). The discrepancy between these numbers is due to a mismatch between the year they were found in the AIS data and the year in which they were detected in tournaments.

**Fig. 1.**
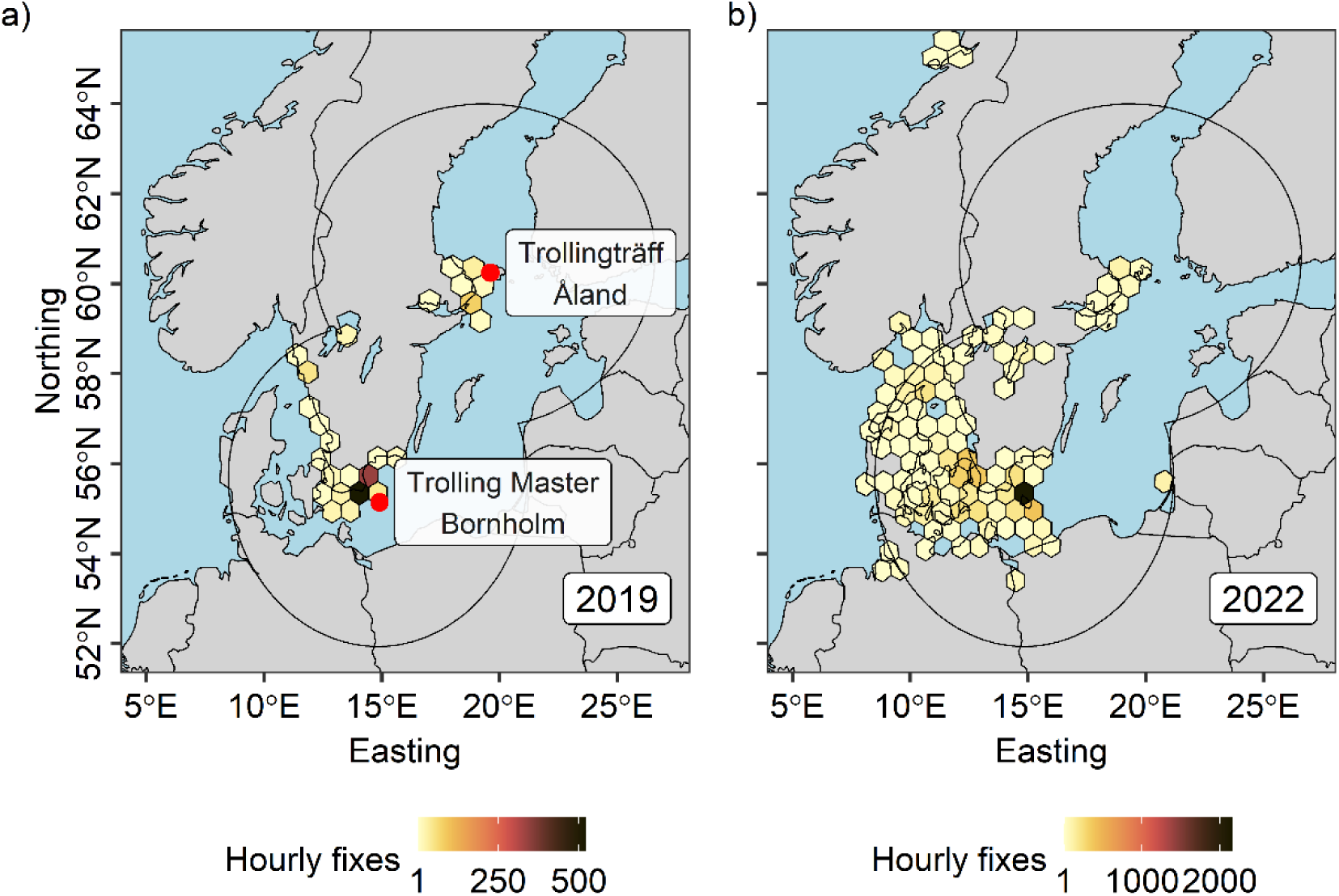
Illustrates the average hourly positions (telemetry fixes) of trolling boat skippers based on automatic identification system (AIS) data. Panel a) displays the positions of 24 skippers for the year 2019, while panel b) shows the positions of 143 skippers for the year 2022, with each skipper represented by 50 randomly sampled fixes. In panel a), the locations of the two tournaments, Trolling Master Bornholm and Trollingträff Åland, are indicated. Notably, in 2019 and 2022, 43% and 30% of the total number of fixes, respectively, fall within the grid cell with the highest number of registrations, situated near the island of Bornholm. In the spatial capture-recapture (SCR) model, 23 of the skippers depicted in panel a) in 2019 were detected in a tournament during session 1 (years 2014-2020), while 111 of the skippers depicted in panel b) in 2022 were detected in a tournament during session 2 (years 2021-2023). The circular area extending from the tournament locations represents the extent of the state space, which is the area used for estimating the population. Telemetry fixes outside this buffer were removed from the SCR analyses to avoid truncated fixes.

A total of 1,093 Baltic Sea salmon trolling boat skippers were detected in tournament lists from 2014 to 2023. From 2014-2020, 937 unique skippers were detected, while from 2021-2023, 458 unique skippers were detected. There were 14 tournaments with data during 2014-2023, with six in Åland and eight in Bornholm. The COVID-19 pandemic caused the cancellation of the tournaments in Bornholm in 2020 and 2021, and no data was collected from Åland from 2014 to 2017 (Table 1).

**Table 1:**
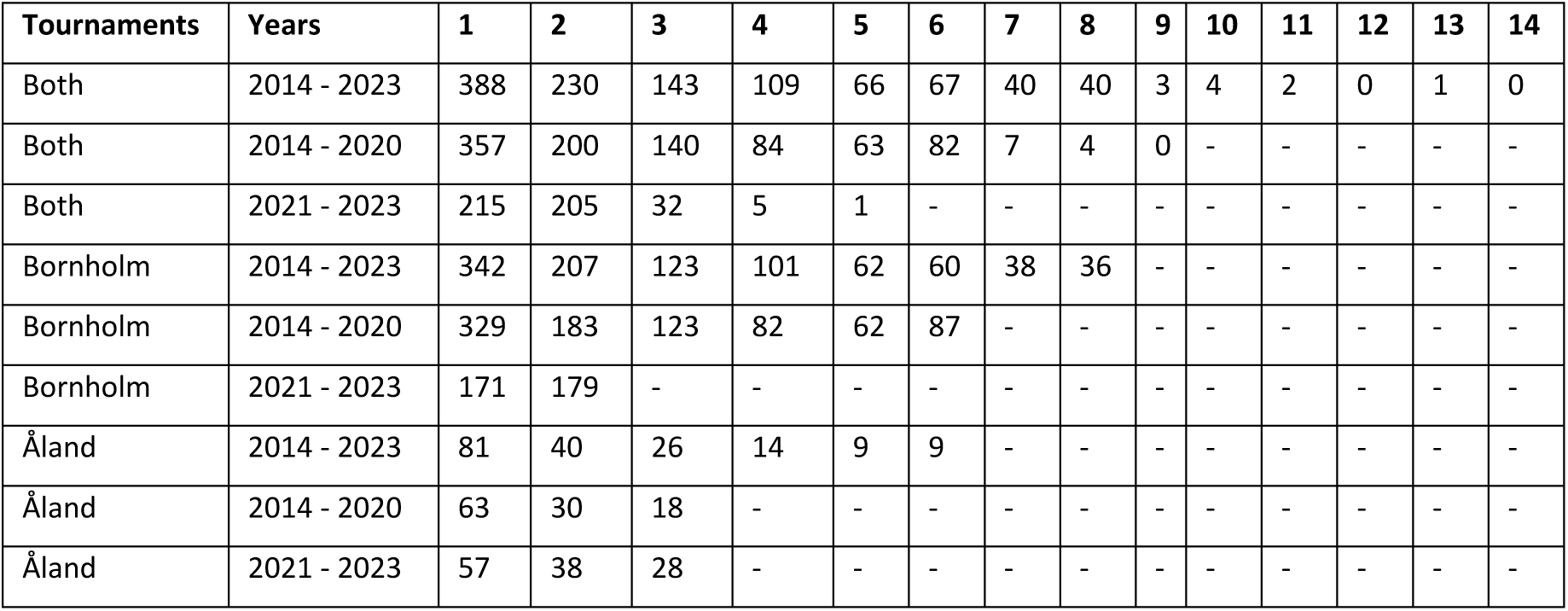
Summary of skipper detection frequencies in tournaments. This table presents the frequencies of the number of detections per skipper in both the Trolling Master Bornholm and the Trollingträff Åland, as well as in each tournament separately. The “Years” column show 2014-2023 corresponding to both session 1 and 2 in the models, where 2014-2020 is designated as session 1, and 2021-2023 is session 2. Trolling Master Bornholm was closed due to the COVID-19 pandemic in 2020 and 2021, resulting in no skippers being detected during those years. Additionally there is no data from Trollingträff Åland for the years 2014-2017. Note that in most rows, the number of skippers that participated at least twice is greater than the number who only participated once.

| Tournaments | Years | 1 | 2 | 3 | 4 | 5 | 6 | 7 | 8 | 9 | 10 | 11 | 12 | 13 | 14 |
| --- | --- | --- | --- | --- | --- | --- | --- | --- | --- | --- | --- | --- | --- | --- | --- |
| Both | 2014 - 2023 | 388 | 230 | 143 | 109 | 66 | 67 | 40 | 40 | 3 | 4 | 2 | 0 | 1 | 0 |
| Both | 2014 - 2020 | 357 | 200 | 140 | 84 | 63 | 82 | 7 | 4 | 0 | - | - | - | - | - |
| Both | 2021 - 2023 | 215 | 205 | 32 | 5 | 1 | - | - | - | - | - | - | - | - | - |
| Bornholm | 2014 - 2023 | 342 | 207 | 123 | 101 | 62 | 60 | 38 | 36 | - | - | - | - | - | - |
| Bornholm | 2014 - 2020 | 329 | 183 | 123 | 82 | 62 | 87 | - | - | - | - | - | - | - | - |
| Bornholm | 2021 - 2023 | 171 | 179 | - | - | - | - | - | - | - | - | - | - | - | - |
| Åland | 2014 - 2023 | 81 | 40 | 26 | 14 | 9 | 9 | - | - | - | - | - | - | - | - |
| Åland | 2014 - 2020 | 63 | 30 | 18 | - | - | - | - | - | - | - | - | - | - | - |
| Åland | 2021 - 2023 | 57 | 38 | 28 | - | - | - | - | - | - | - | - | - | - | - |

In Bornholm, Denmark, most skippers were Danish residents, with Swedes being the second most common residency (Fig. 2a). In Åland, Finland, most skippers were Finnish residents, followed by Swedes as the second most common residency (Fig. 2b). Across both tournaments, Danish residency was the most common, followed by Swedish (Fig. 2c).

**Fig. 2.**
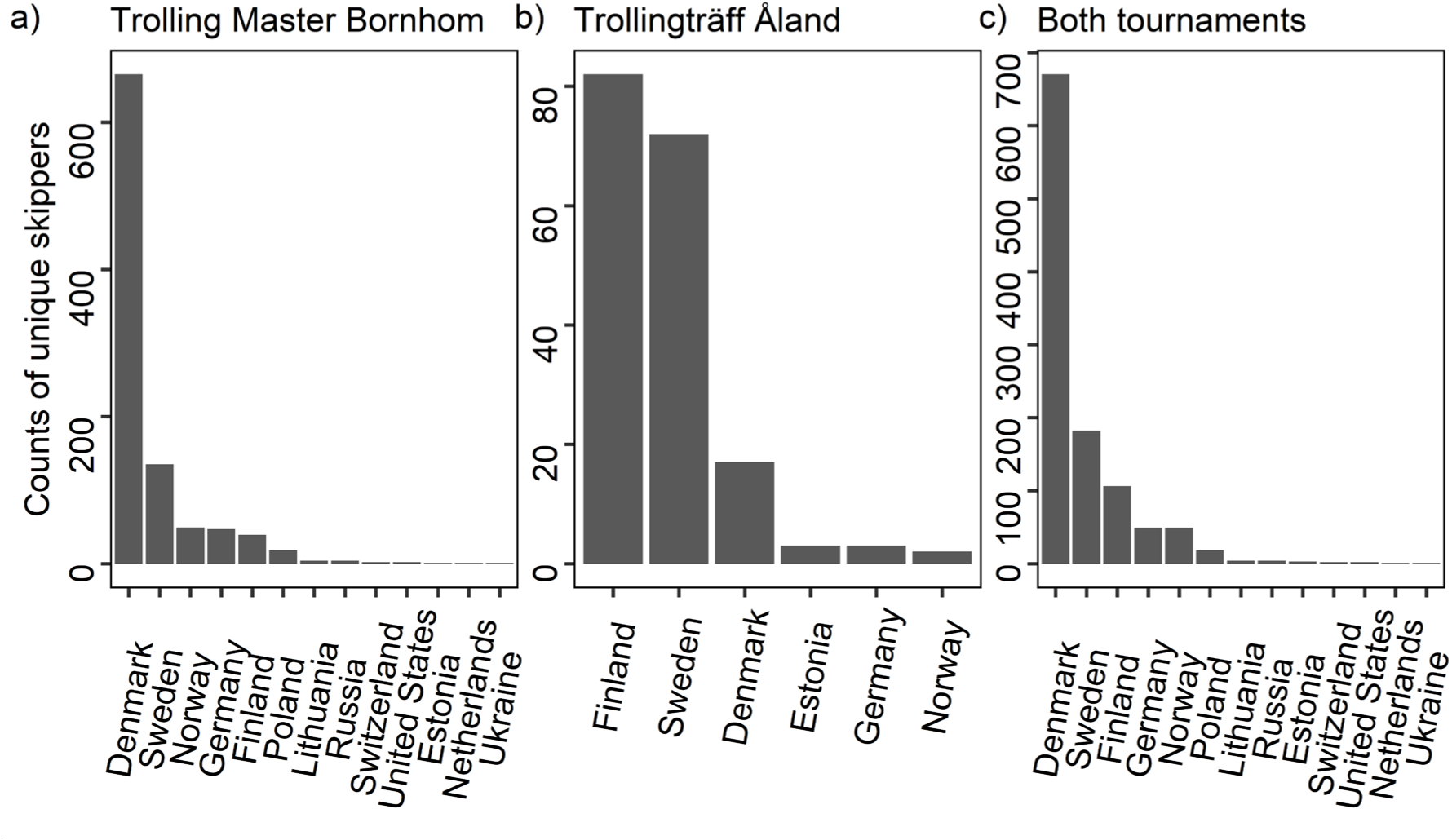
Depicts the unique participants by country of residence in a) Trolling Master Bornholm (Denmark), b) Trollingträff Åland (Finland), and c) both tournaments, spanning the years 2014-2023.

The top model received overwhelming support based on AIC ranking, with ΔAIC = 42.3 (Table 2). Predictions from this model indicated a much lower density of Baltic Sea salmon trolling boat skippers in 2021-2023 compared to 2014-2020, with estimates of 2,604 (95% CI: 2,273-2,983) and 5,343 (4,622-6,178), respectively (Table 2). This corresponds to a 51% decrease between sessions. The population extent is illustrated in Figure 1ab and Figure 3a.

**Fig. 3.**
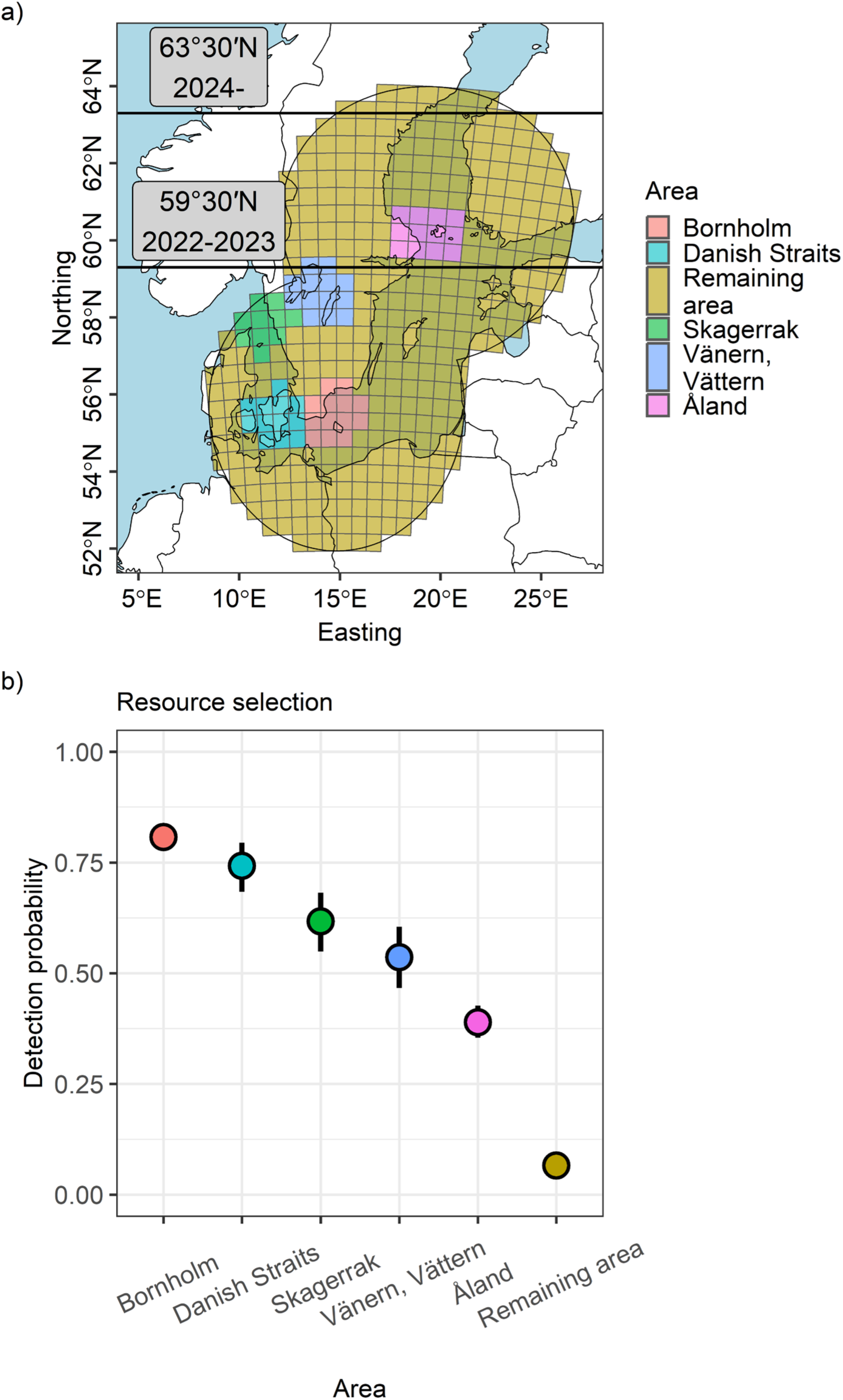
Panel a) illustrates the six different areas within the resource selection data frame. The five areas—Bornholm, Danish straits, Skagerrak, Vänern and Vättern, and Åland—are well-known and popular trolling areas for recreational fishing. The ‘Remaining area’ essentially denotes the region not designated as a trolling area; however, trolling fishing may still be conducted here. The circular area extending from the tournament locations in panel a) represents the extent of the state space, i.e., the area in which the population is estimated. The lines in panel a) labelled 59°30’N (expired) and 63°30’N (current) indicates a change in fishing regulations. North of the line, there are no harvest restrictions on farmed salmon between May 1^st^ to August 31^st^, within 4 nautical miles from the baseline, although wild salmon must still be released. South of these lines, all wild salmon has to be released and only one farmed fish is allowed to be kept per person and day. Panel b) displays the capture probability predictions from the top model (based on AIC ranking), where areas are color-coded and plotted on the x-axis. A higher detection probability means that the tiles have greater importance as a resource for trolling anglers. Error bars represent the 95% confidence interval. For each area predictions were made at a distance of zero from trap to home range center, i.e., *distances* (*x*_j_, *s*_*i*_)^2^, = 0, and averaged across the remaining covariates.

**Table 2:** Model selection analysis for population estimates of Baltic Sea salmon trolling boat skippers. This table presents the model selection results for the top ten models out of 64. The “Density” column displays the covariate ‘session’, representing the two different time periods: 2014-2020 and 2021-2023. The “Detection” column indicates the resource selection covariate ‘area’, the ‘session’ covariate related to time periods, the behavioral covariate ‘b’, which estimates whether previous participation in tournaments affects future participation, and nation of residency allowing detection to vary between the two groups: Danish residents and residents of all other nations. The sigma model is not included in the table as all models in the top ten incorporate the nation of residency covariate. A ‘1’ indicates models with only an intercept. ℒ represents the model log likelihood, np denotes the number of parameters, AIC shows the Akaike Information Criterion used for ranking the models, ΔAIC displays the AIC ranking with the lowest AIC model on top, AICω represents the model weights based on AIC, and AICω+ indicates the cumulative model weight. The columns Ň1 and Ň2 provide the estimates and 95% confidence intervals of the population size of Baltic Sea salmon trolling boat skippers from each model for the two respective sessions.

| Density | Detection | $\mathcal{L}$ | np | AIC | $\Delta AIC$ | AICω | AIC$\omega+$ | $\hat{N}1$ | $\hat{N}2$ |
| --- | --- | --- | --- | --- | --- | --- | --- | --- | --- |
| D(session) | p(area + session<br>+ nation of residency<br>+ b) | 19005.64 | 14 | 38039.28 | 0 | 1 | 1 | 5343<br>(4622-6178) | 2604<br>(2273-2983) |
| D(session) | p(area<br>+ nation of residency + b) | 19027.79 | 13 | 38081.57 | 42.3 | 0 | 1 | 5095<br>(4570-5680) | 3411<br>(2985-3898) |
| D(session) | p(area<br>+ session + nation of residency) | 19043.67 | 13 | 38113.35 | 74.07 | 0 | 1 | 4549<br>(4152-4985) | 2187<br>(1955-2446) |
| D(1) | p(area<br>+ nation of residency + b) | 19052.7 | 12 | 38129.4 | 90.13 | 0 | 1 | 4615<br>(4250-5011) | 4615<br>(4250-5011) |
| D(1) | p(area + session<br>+ nation of residency<br>+ b) | 19052.68 | 13 | 38131.36 | 92.08 | 0 | 1 | 4606<br>(4242-5000) | 4606<br>(4242-5000) |
| D(session) | p(area<br>+ nation of residency) | 19065.76 | 12 | 38155.52 | 116.25 | 0 | 1 | 4339<br>(3971-4742) | 2758<br>(2463-3089) |
| D(session) | p(area<br>+ session + b) | 19066.08 | 13 | 38158.16 | 118.89 | 0 | 1 | 5032<br>(4473-5659) | 1992<br>(1509-2629) |
| D(1) | p(area<br>+ nation of residency) | 19098.57 | 11 | 38219.13 | 179.85 | 0 | 1 | 3750<br>(3587-3922) | 3750<br>(3587-3922) |
| D(1) | p(area<br>+ session + nation of residency) | 19098.62 | 12 | 38221.24 | 181.96 | 0 | 1 | 3731<br>(3441-4045) | 3731<br>(3441-4045) |
| D(session) | p(area<br>+ session) | 19109.4 | 12 | 38242.79 | 203.51 | 0 | 1 | 4167<br>(3513-4941) | 1653<br>(1432-1908) |

Detection probability was influenced by all of the four included covariates: area, behavior, session, and residency (Table 3). Resource selection, estimated as detection probability, was the greatest around the island of Bornholm in the Baltic Sea and the lowest in the area designated as ‘remaining area’, i.e., an area not well-known for trolling fishing (Fig. 3ab). The behavioral response showed that detection probability of trolling skippers that previously had participated in a tournament was greater than for those that not yet had participated (Fig. 4a). The inclusion of a session covariate revealed that the detection probability was greater during 2021-2023 than 2014-2020 (Fig. 4b). Finally, Danish residents had a lower detection probability than residents of other nations (Fig. 4c).

**Fig. 4.**
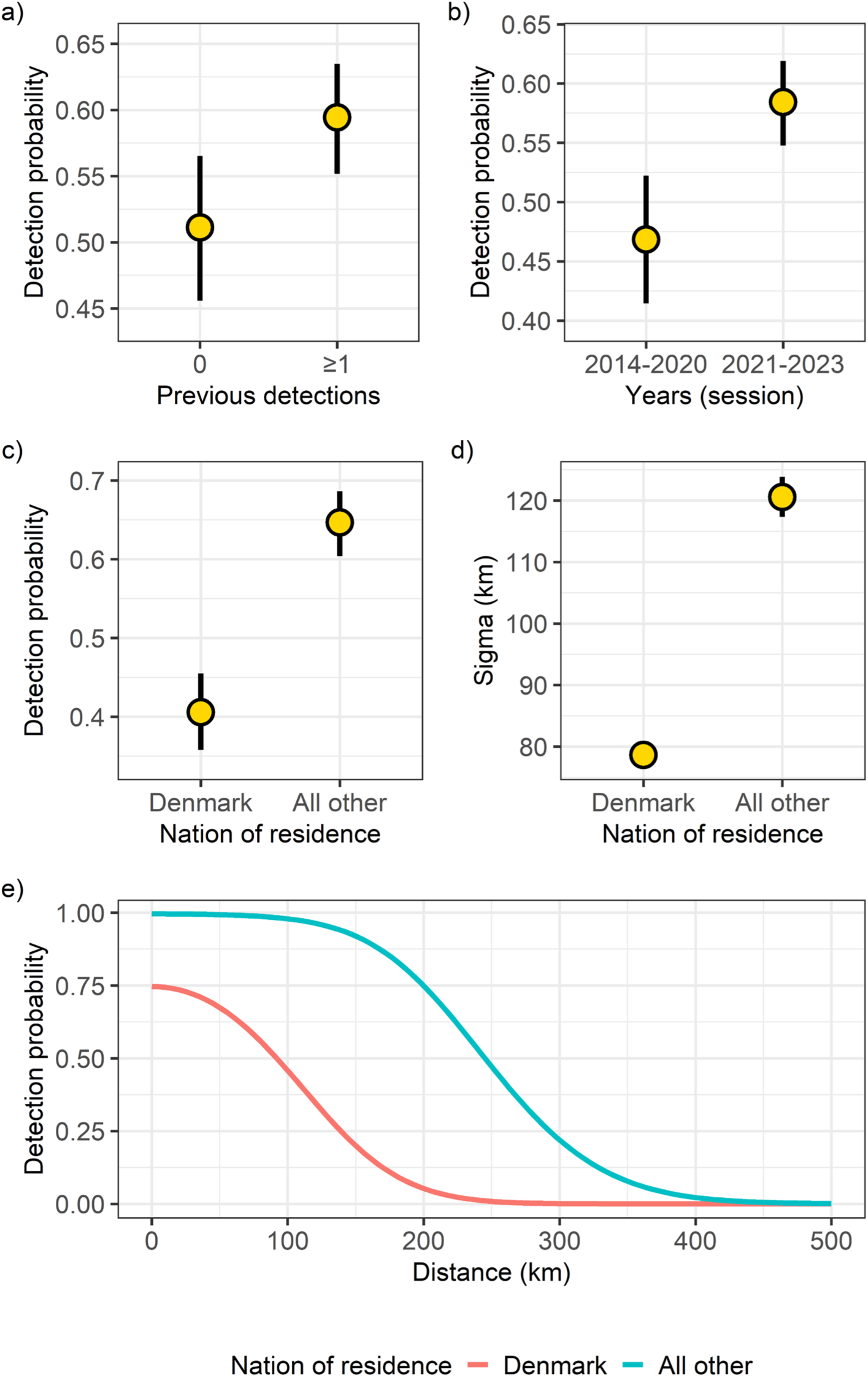
Displays three panels, a) to c), of detection probability predictions and two panels of movement predictions, d) and e), from the top model (based on AIC ranking). In panel a) the two behavioral groups of zero detections and at least one detection are plotted on the x-axis. Panel b) shows the difference in detection probability in the two sessions, with a greater probability in recent years. In panel c), Danish residents, the most common nation of residence for trolling anglers in the Baltic Sea, are compared to residents from all other nations in terms of detection probability. Similarly, in panel d) Danish residents are compared to residents from all other nations regarding the movement parameter sigma. In panel e), the detection probability and sigma are combined to calculate detection probability across the entire home range of Danish residents and the group of residents from all other nations. This is the half-normal detection model described in the methods, where the p0 intercept and sigma were taken from the top model. Error bars in a) to d) represents the 95% confidence interval. In panel a) to d) predictions are made at a distance of zero from the trap to the home range center, i.e., *distances* (*x*_j_, *s*_*i*_)^2^, = 0, and are averaged across the six areas Bornholm, Danish Straits, Vänern and Vättern, Remaining area, Skagerrak, and Åland (Fig 3a).

**Table 3:**
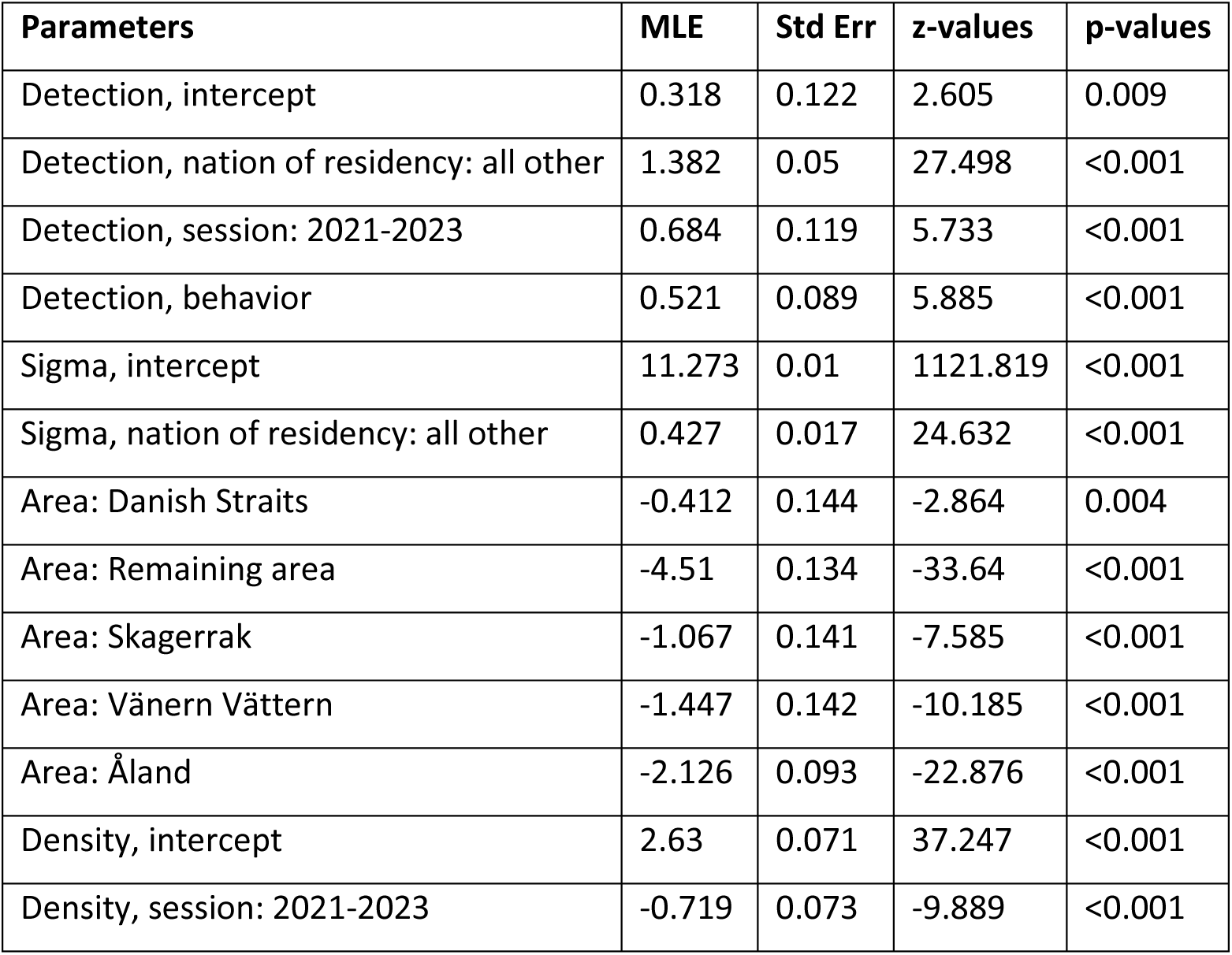
Parameter estimates and statistics for the top model. This table presents the maximum likelihood estimates (MLE), standard errors (Std Err), z-values, and p-values of the top model, as determined by AIC rankings (refer to Table 2). The parameter p0 represents the complementary log-log scale detection probability. The p0 intercept includes the reference levels: Danish residents, the session 2014-2020, the behavior group with zero previous detections, and the Bornholm area. The nation of residency parameter indicates that individuals with a residency other than Danish have a greater detection probability. The session parameter shows that detection probability was greater during 2021-2023 than during 2014-2020. Behavior denotes the effect on detection probability based on previous captures, with previously detected individuals having a greater probability of being detected than the baseline. The area Bornholm had a greater detection probability than all other areas, indicating its importance for trolling skippers. Sigma indicates the log scale movement parameter, with its baseline being Danish residents, who have smaller home ranges than individuals from with other residencies, as reflected in the sigma nationality parameter. Finally, the log scale density parameter, with the intercept corresponding to the years 2014-2020, shows that there were fewer anglers in recent years (2021-2023).

The movement model sigma showed that skippers residing in Denmark moved over shorter distances (Fig. 4d) and had smaller home ranges than skippers from other nations (Fig. 4e).

## Discussion

A key result from this study is a significant decline of approximately 51% in the number of salmon (*Salmo salar*) trolling skippers in the Baltic Sea from the years 2014-2020 to 2021-2023. Due to the data structure with time frames of different length, seven years versus three years, both time frames were estimated with independent detection probabilities. The years 2021-2023 had, as one can expect, fewer detections, but more importantly also a greater detection probability. As the population size essentially is estimated as the ratio between the number of detected individuals and the detection probability, this led to the estimated decrease in the population size of skippers. This decline in numbers is likely due to an increased regulation of the salmon trolling fishery, which no longer allows the harvest of wild salmon in the Baltic Proper and Bothnian Sea, and in most cases, only permits one farmed salmon to be harvested per day and person (ICES, 2023), where the Bothnian Bay north of latitude 63°30′N is the exception. These changing regulations likely reduced angler satisfaction, and together with changes in the global economy in 2022, with rising fuel prices and inflation, this may have affected the population negatively. Notably, in 2024, the Trolling Master Bornholm was canceled because the organizers did not know how to incorporate these new regulations into the competition. Furthermore, Trollingträff Åland only had 17 teams participating in 2024 in contrast to the average of 66 teams participating in the years 2018-2023.

Further, to maintain and support fisheries an average of 4.4 million salmon smolt were released into the Baltic Sea from hatcheries between 2018 to 2022 (ICES, 2023). However, due to COVID restrictions at the hatcheries, out of the 4.6 million released smolt in 2021, only 1.9 million were fin-clipped, compared to the annual average of 3.3 million (ICES, 2023). Because fin-clipping indicates that the salmon is farmed and can be harvested, the number of harvestable salmon in the Baltic Sea is likely to decrease in the coming years. A gradual decrease of fin-clipped salmon is anticipated in 2023 as this cohort will have spent one winter at sea and begins to be targeted by troll fishers (see Figure 2 in Karlsson and Karlström, 1994). Stronger effects of this cohort are expected in 2024 and 2025. A reduction in the number of fin-clipped salmon released into the Baltic Sea is likely to further increase angler dissatisfaction. As a result, the reduced availability of harvestable salmon is likely to lead to a decline in participation in this fishery in general, and in trolling tournaments in particular (Birdsong et al., 2021).

Resource selection was estimated for each pixel of the state-space by integrating the trolling boats’ Automatic Identification System (AIS) data as telemetry fixes to the SCR model. Pixels with greater estimated detection probability are more important as resources for the population (Royle et al., 2013b; Linden et al., 2018), suggesting that Bornholm is the most important area for this population. This is an expected finding as the Trolling Master Bornholm attracts hundreds of anglers yearly, and that the AIS data showed that more than 30% of all trolling boat registrations fall near the island of Bornholm. The next most important area is the Danish Straits, a highly populated region near major cities. In contrast to Bornholm, most of this area is not used for trolling for salmon, so other species are in focus here, such as sea-run brown trout (*Salmo trutta*), which are common nearshore along these islands (Thorstad et al., 2016; Kristensen et al., 2019). The third most important area was Skagerrak, where focus is also on species other than salmon, such as Bluefin tuna (*Thunnus thynnus*). In Skagerrak recreational anglers have been involved in research projects to capture and tag individuals with electronic pop-up satellite archival tags (Birnie-Gauvin et al., 2020; Aarestrup et al., 2022). The Bluefin tuna migrate to Skagerrak and Kattegat from their spawning grounds in the Mexican Gulf and Mediterranean Sea (Aarestrup et al., 2022) where they have recently shown signs of recovery due to conservation efforts (ICCAT, 2006; Aarestrup et al., 2022). The trolling areas of least importance were the Swedish Great Lakes, Vänern and Vättern, which host a popular salmon troll fishery (Andersson et al., 2020), and the area off the Åland archipelago in the Northern Baltic Proper. The remaining area of the population extent had comparatively very low importance for trolling skippers.

The results from the resource selection analysis show that areas outside of the Baltic Sea, and species other than salmon, are also important for this population of skippers. Meaning that these skippers establish a link between different populations of fish. For example, the same population of skippers targets a wild population of Baltic salmon native to the Purtse River in Estonia (Kesler et al., 2011), and a landlocked wild population of salmon in the Gullspång River, native to Lake Vänern in Sweden (Piccolo et al., 2012). This indicates that restrictions on one stock may have unforeseen cascading effects on another if skippers shift areas depending on regulations. This fact is important for fisheries managers to be aware of, and restrictions should not be harder than necessary as this likely intensify fishing in other areas. Furthermore, the relaxed regulations in the Northern Baltic Sea (Bothnian Bay) likely have little effect on the northern salmon populations as this area is relatively unimportant for trolling skippers and these populations are relatively viable. Instead, the regulations may relieve sensitive southern populations in more attractive areas from high fishing pressure. This finding agrees well with the recently implemented area specific restrictions.

The model results showed that trolling boat skippers are a heterogeneous group in terms of tournament participation. Skippers who had previously participated in a tournament showed a higher detection probability than those who had not, indicating a group of highly dedicated skippers that participate more frequently. This pattern is referred to as “trap-happiness” (a positive response), or “trap-shyness” (a negative response) within the capture-recapture framework (Chapter 7 in Royle et al., 2013a). In datasets where trap happy effects are present, the null model overestimate detection probability because individuals who are never observed have lower detection probability than those with previous detections. Consequently, if this parameter is excluded, the population size will be underestimated. Although there were fewer skippers during 2021-2023 they were more likely to participate in tournaments, which further emphasizes the strong dedication of some skippers to tournament angling, while also highlighting the recent decline in less dedicated skippers. These results suggest that, for most skippers, tournament participation is a positive experience.

The binary detection covariate ‘nation of residency’ estimated that skippers residing in Denmark had a lower detection probability than skippers residing in other countries, which may be due to their proximity to the Trolling Master Bornholm, allowing for more spontaneous participation. Thus, Danish residents may participate just occasionally. Whereas skippers residing in other nations, who must travel farther, may have a greater devotion to participate in tournaments and, therefore, a higher detection probability. This theory is further supported by the movement scale parameter sigma. Danish residents travelled much shorter distances than those from other nations, which could be caused by their proximity to important fishing grounds in general, and to the Trolling Master Bornholm specifically. A skipper living in Denmark has a home range radius of approximately 250 km, compared to 450 km for skippers residing in other nations. Thanks to the numerous AIS detections, the movements of skippers could be estimated with great precision, with a ratio between estimate and standard error of approximately 1122 for the sigma intercept.

### Conclusions, limitations, and future directions

The well-developed SCR framework includes models that can estimate many variables necessary for describing human populations. Among others variables, both skipper behavior and their movements are incorporated in the detection probability function, resulting in more realistic estimates of population density and extent. Naturally, not every aspect of the skipper population can be included in the models, and assumptions about the population must be understood in part through the included covariates. Overly complex models, which include too many assumptions about the population, are often data-deficient, do not generalize well, and are more likely to incorporate less significant parameters that may not be relevant over time (Chapter 4 in Royle et al., 2013a).

A limitation of this type of study is the sensitive nature of working with personal data, which may affect the future scalability of modeling tournament participation as encounter histories.

Furthermore, for this study in particular, the low spatial resolution relying on only two tournament locations was mitigated by integrating AIS data, which allowed for estimating resource selection and movements. However, only two sampling locations does not allow to estimate variation in density, which is currently assumed to be uniformly distributed across the state space, although most likely skipper density is higher near populated areas and important fishing grounds.

In a fishery that heavily restricts harvest but still allows the catching of salmon (Thorstad et al., 2003), estimating catch-and-release mortality becomes increasingly important. Currently there are no published studies on catch-and-release survival of salmon in the Baltic Sea troll fishery, though many studies address survival in rivers (Keefe et al., 2022; Van Leeuwen et al., 2021; Van Leeuwen et al., 2023). Survival at sea is an important future topic to better quantify the effects of recreational fishing on Baltic salmon populations.

## Data availability

An anonymized version of the spatial capture-recapture data and the script used to perform the models will be made available during peer review

## Declaration of competing interests

None to declare

## Funding

AIS data was purchased by funds from the Data Collection Framework (DCF) which provides a common framework for the EU community to collect, manage, and share data within the fisheries sector.

## Acknowledgements

I am grateful for the constructive comments from colleagues during the presentation of this work at the department and at the ICES WGBAST meeting in Gävle, Sweden, in 2024. I would also like to thank Jon Brommer for organizing a great workshop held at the University of Turku, Finland, in 2016, where Andrew Royle and Angela Fuller presented their work on spatially explicit capture-recapture modeling.

## Notes

### Competing Interest Statement

The authors have declared no competing interest.

## References

Aarestrup, K., Baktoft, H., Birnie-Gauvin, K., Sundelöf, A., Cardinale, M., Quilez-Badia, G., Onandia, I., Casini, M., Nielsen, E.E., Koed, A. & Alemany, F. (2022). First tagging data on large Atlantic bluefin tuna returning to Nordic waters suggest repeated behaviour and skipped spawning. Scientific reports, 12(1), p.11772. 10.1038/s41598-022-15819-x

Andersson, A., Greenberg, L. A., Bergman, E., Su, Z., Andersson, M., & Piccolo, J. J. (2020). Recreational trolling effort and catch of Atlantic salmon and brown trout in Vänern, the EU’s largest lake. Fisheries research, 227, 105548. 10.1016/j.fishres.2020.105548

Birdsong, M., Hunt, L. M., & Arlinghaus, R. (2021). Recreational angler satisfaction: What drives it?. Fish and Fisheries, 22(4), 682–706. 10.1111/faf.12545

Birnie-Gauvin, K., MacKenzie, B. R., & Aarestrup, K. (2020). Electronic tagging of Atlantic bluefin tunas in Scandinavian waters 2018. Collect. Vol. Sci. Pap. ICCAT, 76(2), 667-672. https://www.iccat.int/Documents/CVSP/CV076_2019/n_2/CV076020667.pdf

Borchers, D. L., & Efford, M. G. (2008). Spatially explicit maximum likelihood methods for capture– recapture studies. Biometrics, 64(2), 377–385. 10.1111/j.1541-0420.2007.00927.x

Chao, A., Tsay, P. K., Lin, S. H., Shau, W. Y., & Chao, D. Y. (2001). The applications of capture-recapture models to epidemiological data. Statistics in medicine, 20(20), 3123–3157. 10.1002/sim.996

Efford, M. (2004). Density estimation in live-trapping studies. Oikos, 106(3), 598–610. 10.1111/j.0030-1299.2004.13043.x

Fuller, A. K., Sutherland, C. S., Royle, J. A., & Hare, M. P. (2016). Estimating population density and connectivity of American mink using spatial capture–recapture. Ecological Applications, 26(4), 1125–1135. 10.1890/15-0315

Hartill, B. W., Taylor, S. M., Keller, K., & Weltersbach, M. S. (2020). Digital camera monitoring of recreational fishing effort: applications and challenges. Fish and Fisheries, 21(1), 204–215. 10.1111/faf.12413

HELCOM. (2011). Salmon and Sea Trout Populations and Rivers in the Baltic Sea–HELCOM assessment of salmon (*Salmo salar*) and sea trout (*Salmo trutta*) populations and habitats in rivers flowing to the Baltic Sea. In Baltic Sea Environment Proceedings (Vol. 126, p. 79).

ICCAT (2006). Recommendation by ICCAT to Establish a Multi-annual Recovery Plan for Bluefin Tuna in the Eastern Atlantic and Mediterranean. Recommendation 06–05. https://www.iccat.int/Documents/Recs/compendiopdf-e/2006-05-e.pdf

ICES (2021). Baltic Salmon and Trout Assessment Working Group (WGBAST). ICES Scientific Reports. Report. 10.17895/ices.pub.7925

ICES (2023). Baltic Salmon and Trout Assessment Working Group (WGBAST). ICES Scientific Reports. Report. 10.17895/ices.pub.22800983.v1

Jacobson, P., Gårdmark, A., & Huss, M. (2020). Population and size-specific distribution of Atlantic salmon *Salmo salar* in the Baltic Sea over five decades. Journal of Fish Biology, 96(2), 408–417. 10.1111/jfb.14213

Jones, D., Dahlgren, E., Jacobson, P., & Karlson, A. M. (2022). Determining Baltic salmon foraging areas at sea using stable isotopes in scales—a tool for understanding health syndromes. ICES Journal of Marine Science, 79(1), 158–168. 10.1093/icesjms/fsab250

Kallio-Nyberg, I., & Ikonen, E. (1992). Migration pattern of two salmon stocks in the Baltic Sea. ICES Journal of Marine Science, 49(2), 191–198. 10.1093/icesjms/49.2.191

Karlsson, L., & Karlström, Ö. (1994). The Baltic salmon (*Salmo salar* L.): its history, present situation and future. Dana, 10, 61–85.

Keefe, D., Young, M., Van Leeuwen, T. E., & Adams, B. (2022). Long-term survival of Atlantic salmon following catch and release: Considerations for anglers, scientists and resource managers. Fisheries Management and Ecology, 29(3), 286–297. 10.1111/fme.12533

Kéry, M., & Royle, J. A. (2021). Applied hierarchical modeling in ecology. Volume 2, Dynamic and advanced models: Analysis of distribution, abundance and species richness in R and BUGS. Academic Press.

Kesler, M., Kangur, M., & Vetemaa, M. (2011). Natural re-establishment of Atlantic salmon reproduction and the fish community in the previously heavily polluted River Purtse, Baltic Sea. Ecology of freshwater fish, 20(3), 472–477. 10.1111/j.1600-0633.2010.00483.x

Klemetsen, A., Amundsen, P. A., Dempson, J. B., Jonsson, B., Jonsson, N., O’connell, M. F., & Mortensen, E. (2003). Atlantic salmon *Salmo salar* L., brown trout Salmo trutta L. and Arctic charr Salvelinus alpinus (L.): a review of aspects of their life histories. Ecology of freshwater fish, 12(1), 1–59. 10.1034/j.1600-0633.2003.00010.x

Kristensen, M.L., Pedersen, M.W., Thygesen, U.H., del Villar-Guerra, D., Baktoft, H. & Aarestrup, K., (2019). Migration routes and habitat use of a highly adaptable salmonid (sea trout, Salmo trutta) in a complex marine area. Animal biotelemetry, 7, pp.1–13. 10.1186/s40317-019-0185-3

Kärgenberg, E., Thorstad, E. B., Järvekülg, R., Sandlund, O. T., Saadre, E., Økland, F., Thalfeldt, M., & Tambets, M. (2019). Behaviour and mortality of downstream migrating Atlantic salmon smolts at a small power station with multiple migration routes. Fisheries Management and Ecology, 27(1), 32–40. 10.1111/fme.12382

Linden, D. W., Sirén, A. P., & Pekins, P. J. (2018). Integrating telemetry data into spatial capture– recapture modifies inferences on multi-scale resource selection. Ecosphere, 9(4), e02203. 10.1002/ecs2.2203

Miettinen, A., Palm, S., Dannewitz, J., Lind, E., Primmer, C. R., Romakkaniemi, A., Östergren, J., & Pritchard, V. L. (2021). A large wild salmon stock shows genetic and life history differentiation within, but not between, rivers. Conservation Genetics, 22(1), 35–51. 10.1007/s10592-020-01317-y

Otis, D. L., Burnham, K. P., White, G. C., & Anderson, D. R. (1978). Statistical inference from capture data on closed animal populations. Wildlife monographs, (62), 3-135. http://www.jstor.org/stable/3830650

Palm, S., Dannewitz, J., Järvi, T., Koljonen, M. L., Prestegaard, T., & Olsén, K. H. (2008). No indications of Atlantic salmon (*Salmo salar*) shoaling with kin in the Baltic Sea. Canadian Journal of Fisheries and Aquatic Sciences, 65(8), 1738–1748. 10.1139/F08-088

Piccolo, J. J., Norrgård, J. R., Greenberg, L. A., Schmitz, M., & Bergman, E. (2012). Conservation of endemic landlocked salmonids in regulated rivers: a case-study from Lake Vänern, Sweden. Fish and Fisheries, 13(4), 418–433. 10.1111/j.1467-2979.2011.00437.x

Royle, J. A., Chandler, R. B., Sollmann, R., & Gardner, B. (2013a). Spatial capture-recapture. Academic press.

Royle, J. A., Chandler, R. B., Sun, C. C., & Fuller, A. K. (2013b). Integrating resource selection information with spatial capture–recapture. Methods in Ecology and Evolution, 4(6), 520–530. 10.1111/2041-210X.12039

Royle, J. A., Fuller, A. K., & Sutherland, C. (2018). Unifying population and landscape ecology with spatial capture–recapture. Ecography, 41(3), 444–456. 10.1111/ecog.03170

Siira, A., Erkinaro, J., Jounela, P., & Suuronen, P. (2009). Run timing and migration routes of returning Atlantic salmon in the Northern Baltic Sea: implications for fisheries management. Fisheries Management and Ecology, 16(3), 177–190. 10.1111/j.1365-2400.2009.00654.x

Sutherland, C., Royle, J. A., & Linden, D. W. (2019). oSCR: a spatial capture–recapture R package for inference about spatial ecological processes. Ecography, 42(9), 1459–1469. 10.1111/ecog.04551

Thorstad, E. B., Næsje, T. F., Fiske, P., & Finstad, B. (2003). Effects of hook and release on Atlantic salmon in the River Alta, northern Norway. Fisheries Research, 60(2-3), 293–307. 10.1016/S0165-7836(02)00176-5

Thorstad, E.B., Todd, C.D., Uglem, I., Bjørn, P.A., Gargan, P.G., Vollset, K.W., Halttunen, E., Kålås, S., Berg, M. and Finstad, B., (2016). Marine life of the sea trout. Marine Biology, 163, pp.1–19. 10.1007/s00227-016-2820-3

Van Leeuwen, T. E., Dempson, B., Cote, D., Kelly, N. I., & Bates, A. E. (2021). Catchability of Atlantic salmon at high water temperatures: implications for river closure temperature thresholds to catch and release angling. Fisheries Management and Ecology, 28(2), 147–157. 10.1111/fme.12464

Van Leeuwen, T.E., Lehnert, S.J., Breau, C., Fitzsimmons, M., Kelly, N.I., Dempson, J.B., Neville, V.M., Young, M., Keefe, D., Bird, T.J. and Cote, D. (2023). Considerations for water temperature-related fishery closures in recreational Atlantic salmon (*Salmo salar*) catch and release fisheries: A case study from eastern Canada. Reviews in Fisheries Science & Aquaculture, 31(4), pp.598–619. 10.1080/23308249.2023.2242959

Whitlock, R., Mäntyniemi, S., Palm, S., Koljonen, M. L., Dannewitz, J., & Östergren, J. (2018). Integrating genetic analysis of mixed populations with a spatially explicit population dynamics model. Methods in Ecology and Evolution, 9(4), 10.1111/2041-210X.12946

